# Off-Equilibrium Fluctuation-Dissipation Theorem Paves the Way in Alzheimer’s Disease Research

**DOI:** 10.1101/2024.09.15.613131

**Authors:** Gustavo Patow, Juan Monti, Irene Acero-Pousa, Sebastián Idesis, Anira Escrichs, Yonatan Sanz Perl, Petra Ritter, Morten Kringelbach, Gustavo Deco, the Alzheimer’s Disease Neuroimaging Initiative

**Affiliations:** ViRVIG, Universitat de Girona, Girona, Catalonia, Spain; Computational Neuroscience Group, Center for Brain and Cognition, Department of Information and Communication Technologies, Universitat Pompeu Fabra, Barcelona, Catalonia, Spain; Instituto de Física Rosario CONICET-UNR, Laboratorio de Colisiones Atómicas, FCEIA, Universidad Nacional de Rosario, Rosario, Argentina; Cognitive Neuroscience Center (CNC), Universidad de San Andrés, Buenos Aires, Argentina; Berlin Institute of Health at Charité, Charité Universitä tsmedizin Berlin, Robert-Koch-Platz 4, 10117 Berlin, Germany; Department of Psychiatry, University of Oxford, Oxford, UK; Center for Music in the Brain, Department of Clinical Medicine, Aarhus University, Aarhus, Denmark; Institució Catalana de la Recerca i Estudis Avançats (ICREA), Barcelona, Catalonia, Spain

## Abstract

**INTRODUCTION:** Alzheimer’s disease (AD) is a neurodegenerative disorder characterized by progressive cognitive decline. Although traditional methods have provided insights into brain dynamics in AD, they have limitations in capturing non-equilibrium dynamics across disease stages. Recent studies suggest that dynamic functional connectivity in resting-state networks (RSNs) may serve as a biomarker for AD, but the role of deviations from dynamical equilibrium remains underexplored.

**OBJECTIVE:** This study applies the off-equilibrium fluctuation-dissipation theorem (FDT)^1^ to analyze brain dynamics in AD, aiming to compare deviations from equilibrium in healthy controls, patients with mild cognitive impairment (MCI), and those with AD. The goal is to identify potential biomarkers for early AD detection and understand disease progression’s mechanisms.

**METHODS:** We employed a model-free approach based on FDT to analyze functional magnetic resonance imaging (fMRI) data, including healthy controls, MCI patients, and AD patients. Deviations from equilibrium in resting-state brain activity were quantified using fMRI scans. In addition, we performed model-based simulations incorporating Amyloid-Beta (A*β*), tau burdens, and Generative Effective Connectivity (GEC) for each subject.

**RESULTS:** Our findings show that deviations from equilibrium increase during the MCI stage, indicating hyperexcitability, followed by a significant decline in later stages of AD, reflecting neuronal damage. Model-based simulations incorporating A*β* and tau burdens closely replicated these dynamics, especially in AD patients, highlighting their role in disease progression. Healthy controls exhibited lower deviations, while AD patients showed the most significant disruptions in brain dynamics.

**DISCUSSION:** The study demonstrates that the off-equilibrium FDT framework can accurately characterize brain dynamics in AD, providing a potential biomarker for early detection. The increase in non-equilibrium deviations during the MCI stage followed by their decline in AD offers a mechanistic explanation for disease progression. Future research should explore how combining this framework with other dynamic brain measures could further refine diagnostic tools and therapeutic strategies for AD and other neurodegenerative diseases.

## 1 Introduction

Neurodegeneration refers to the progressive loss of structure or function of neurons, often leading to cell death^2^. Among the most common neurodegenerative disorders, Alzheimer’s Disease (AD) is particularly notable for affecting both cortical and subcortical areas of the brain. The disease typically begins in the medial temporal lobe and limbic system and later spreads to most areas of the neocortex^3–5^. Clinically, AD progresses from an asymptomatic phase to a stage characterized by severe cognitive impairments, including memory loss, neuropsychiatric symptoms, and ultimately, a profound decline in all bodily functions. This progression results in a significant loss of quality of life for patients and caregivers and incurs substantial societal costs.

Minor cognitive deficits that minimally affect daily activities are classified as mild cognitive impairment (MCI). Over time, these deficits often extend to other cognitive domains, such as speech and spatial orientation, eventually leading to dementia when daily functioning becomes severely impaired^6^. Neurodegeneration in AD manifests across various levels of neuronal circuitry, from molecular to systemic. Unfortunately, these processes are irreversible, making AD an incurable disease. Current diagnostic methods are inadequate, as evidenced by a 20% misdiagnosis rate^7^, highlighting the urgent need for more accurate diagnostic tools^7^. Research suggests that patients with AD, and to a lesser extent those with MCI, exhibit reduced neuronal connectivity variation during resting-state fMRI, indicating the potential for dynamic functional connectivity (FC) as a biomarker for AD^8^.

Resting-state fMRI has become a valuable tool for exploring the brain’s intrinsic organization of large-scale distributed networks. Recent studies have identified stage-dependent fluctuations in brain activity within several resting-state networks, including the default mode network (DMN), salience network (SN), dorsal attention network (DAN), and limbic networks (LN)^9–17^. However, only a few studies have employed a whole-brain dynamic approach to investigate the impact of AD on brain dynamics and information processing across large-scale networks^18–20^.

On the other hand, the pathological hallmark of AD is the accumulation of misfolded proteins, specifically the extracellular buildup of amyloid-beta (A*β*) plaques and the intraneuronal aggregation of the microtubule-associated protein tau, forming neurofibrillary tangles^21^. Despite advances in developing treatments aimed at reducing A*β* deposition, such as the monoclonal antibodies aducanumab and lecanemab, their effects on cognitive decline remain inconclusive^22^. Although substantial research has focused on understanding AD, many aspects of its pathophysiology, particularly the roles of A*β* and tau, remain poorly understood^2,23^. Regarding brain dysfunction, numerous human and animal studies point to a disruption in the excitation/inhibition (E/I) balance, particularly in the early stages of AD, where neuronal hyperexcitability disrupts cortical function, contributing to cognitive deficits^24,25^. Recent work by Chang et al. demonstrated that tau differentially impacts excitatory and inhibitory neurons, with its removal reducing excitatory neuron activity while simultaneously affecting inhibitory neuron axon initial segments and intrinsic excitability, leading to overall network inhibition^26^. Similarly, Bi et al. hypothesized that A*β* impairs GABAergic function, resulting in synaptic hyperexcitation^27^. Petrache and colleagues corroborated this, showing synaptic hyperexcitation and significant disruptions in the balance of E/I inputs onto principal neurons, alongside a reduction in somatic inhibitory axon terminals^28^. Moreover, Lauterborn et al. recently reported elevated E/I ratios in post-mortem cortical samples, reinforcing the link between E/I imbalance and hyperexcitability in AD^29^. While these studies provide valuable insights into E/I imbalances using animal models and post-mortem human tissue, in vivo investigations in humans are scarce, as neuroimaging methods cannot directly measure the activity of excitatory and inhibitory populations. Most research on whole-brain dynamics has focused on activation patterns without specifically addressing the role of E/I imbalances^30–34^. To better understand the interplay between pathophysiological processes and brain dynamics, biologically plausible models that integrate empirical data and account for the heterogeneity of brain dynamics are needed^35–37^.

Despite advancements in understanding AD’s effects on brain activity, there remain significant gaps in knowledge regarding how disease progression alters brain dynamics. Specifically, existing studies have not fully explored how non-equilibrium dynamics—fundamental to cognitive processes—play a role in AD. While traditional methods have provided insights into static or minimally dynamic aspects of brain function, they fail to capture the full complexity of brain dynamics, particularly in disease progression^38,39^. This gap in understanding limits our ability to accurately characterize brain states and develop reliable biomarkers for early AD detection.

This paper addresses these gaps by applying a novel, model-free approach based on the fluctuation-dissipation theorem (FDT)^1,40^ to analyze non-equilibrium dynamics in AD. Specifically, we propose using the FDT to compare deviations from equilibrium in different stages of AD (MCI and AD) relative to healthy controls. This framework aims to identify distinctive biomarker candidates for early AD detection and provide dynamic correlates of consciousness. This study offers new insights into the mechanisms driving AD progression by advancing our understanding of how non-equilibrium dynamics underpin brain states. It opens avenues for more precise diagnostic and therapeutic strategies. See Figure 1 for an overview of the methods, described in detail below. To shed further light on the possibilities of this framework, we compared these results with the result of applying the framework to the model-based simulations from our previous work^41^, including the Amyloid-Beta (A*β*) and tau burdens, as well as to the inclusion of a personalized Generative Effective Connectivity (GEC) for each subject.

**Figure 1.**
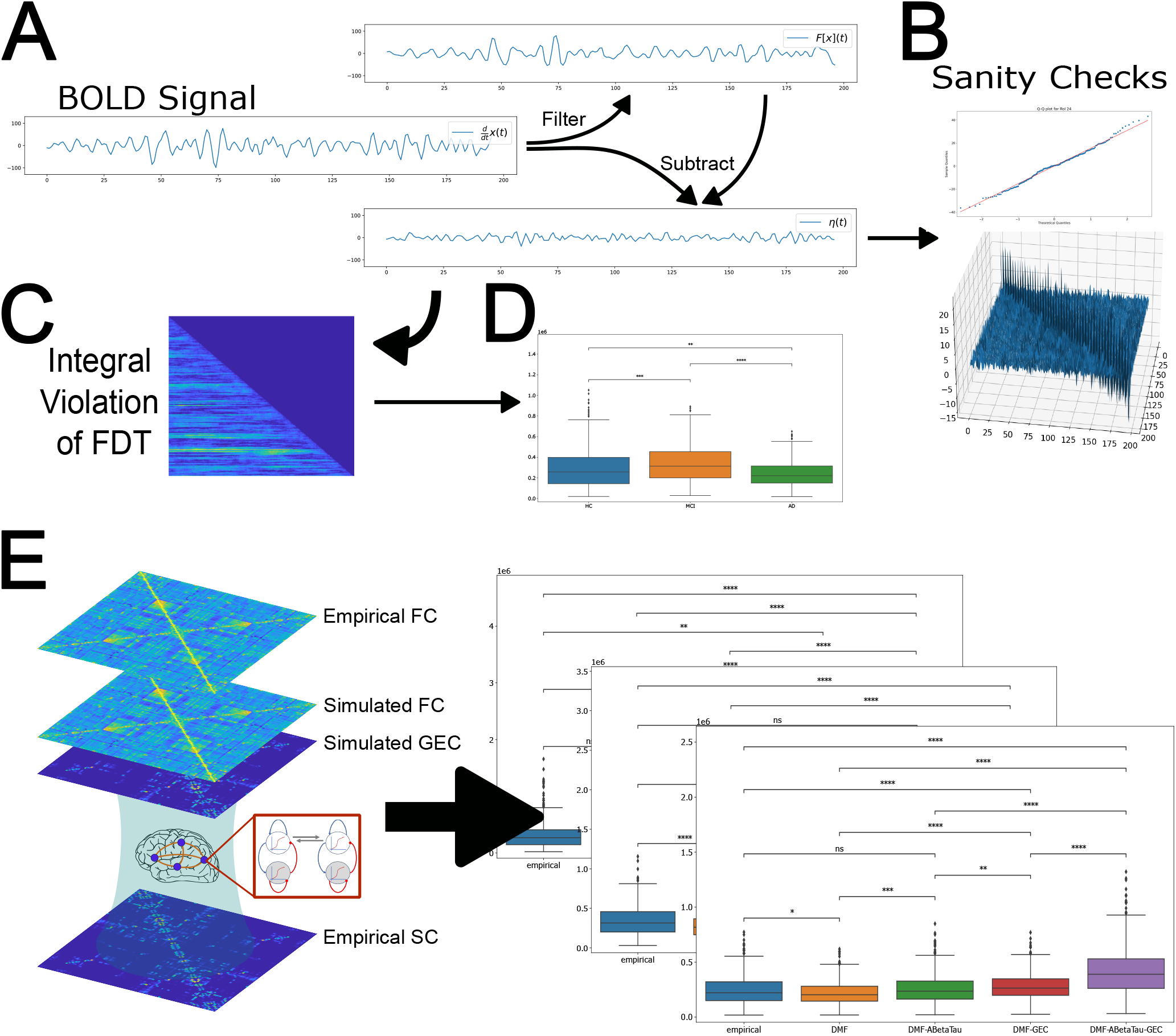
Overview of the Off-Equilibrium FDT Framework and its application for Alzheimer’s Disease. **(A)** The input BOLD signal is numerically derived, and the resulting signal is split into a filtered version and a noise residual *η*(*t*). **(B)** The noise residual is tested for both Gaussianity and the so-called *sanity check*. **(C)** The equilibrium formalism is applied to find the integral violation of the equilibrium, *I*(*t, t*^′^). **(D)** This result is summarized and applied to each condition (HC, MCI, and AD) independently, to verify its analytical power. **(E)** The Model-based pipeline allows comparison of multiple variations of an analytical model to extract information to show the way forward in Alzheimer’s disease research.

## 2 Methods

### 2.1 Participants

We used the ADNI database to gather 36 subjects, subdivided into 17 healthy controls (HC), 9 mild cognitive impairment (MCI) patients, and 10 patients with Alzheimer’s Disease (AD), all taken from ADNI and which are mostly the same participants as those used in previous studies^33,34,41^. See Table 2.

To assess the statistical power of our cohort, we used the G∗Power^42^ software to calculate a two-group Wilcoxon-Mann-Whitney test, with significance level *α* = 0.05 and power 1 − *β* = 0.8. Given our sample size, we can observe that the minimum discernible effect size would be *d* = 1.1. Also, assuming a standard deviation *σ* = 1.65*e*5 (a reasonable assumption given our results below), we get the minimum difference between the deviations from equilibrium must be at least 1.8e5 (in arbitrary units).

### 2.2 Data Acquisition and Processing

The images used in this study were downloaded from ADNI-3, as already mentioned, which used data from Siemens scan. As this data is the same as previously reported^33,41^, we will only briefly describe its preprocessing. The following imaging modalities were included:

- T1 MPRAGE. TE = 2.95– 2.98ms, TR = 2.3 s, matrix and voxel size differ slightly.
- FLAIR. TE differs slightly, TR= 4.8 s, matrix size = 160 × 256 × 256, and voxel size differs slightly.
- DWI (only for 15 HC participants to create an average healthy SC). TE = 56–71ms, TR = 3.4–7.2 s, matrix size = 116 · 116 · 80, voxel size = 2 · 2 · 2, bvals = [0, 1000] or [0, 500, 1000, 2000], bvecs = 49 or 115.
- Siemens Fieldmaps and PET Data (AV-45 for A*β*). The preprocessing of imaging data can be subdivided into structural images, DWI, and PET.

### 2.3 fMRI Pre-Processing

We used FSL FEAT and independent component analysis–based denoising (FSLFIX), following a standard pipeline^33^, which included removal of the first four volumes, rigid-body head motion correction, 3-mm spatial smoothing to improve signal-to-noise ratio, and a high-pass temporal filter of 75s to remove slow drifts to do the pre-processing of resting-state fMRI. Next, we used FSLFIX to denoise the data, an ICA-based (independent component analysis) tool that uses an automated classifier to identify noise-related components for removal from the data. A manually labeled held-out set of 25 individuals scanned with identical imaging parameters was used for training. Next, we removed, from the voxel-wise fMRI time series using ordinary least squares regression, the time courses for noise-labeled components, and 24 head motion parameters.

The resulting denoised functional data were spatially normalized to the NMI152 space (International Consortium for Brain Mapping 152 template in Montreal Neurological Institute) employing the Advanced Normalization Tools (version 2.2.0), in three steps: first, registration of the mean realigned functional scan to the skull-stripped high-resolution anatomical scan via rigid-body registration; second spatial normalization of the anatomical scan to the MNI template via a nonlinear registration; and, finally, normalization of the functional scan to the MNI template using a single transformation matrix that concatenates the transforms generated in the first two steps. Finally, the mean time series for each parcellated region was extracted.

### 2.4 Amyloid-Beta and tau

In Alzheimer’s disease, a defining feature of its progression is the accumulation of amyloid-*β* (A*β*) peptides and tau proteins. This gradual process accumulates, modifies, and aggregates these monomeric forms into larger misfolded structures, eventually forming fibrillar inclusions. This mechanism is believed to be a key driver and initiator of AD. As shown in Figure±6, the levels of these proteins, as assessed by PET imaging, consistently rise throughout the disease’s progression and are often used as biomarkers to gauge the stage of the disease. In our study, the A*β* data was derived from the AV-45 PET scans, which were preprocessed by ADNI, featuring a resolution of 1.5mm cubic voxels and a matrix size of 160 × 160 × 96. The data was normalized to ensure an average voxel intensity of 1, smoothed using a filter specific to the scanner, and then averaged across the Glasser parcellation. For tau, we utilized the preprocessed AV-1451 (Flortaucipir) data from ADNI, following the same acquisition and processing procedures, resulting in a single relative tau value per voxel, which was subsequently averaged according to the corresponding parcellation values.

### 2.5 Deviation from Off-Equilibrium FDT Formalism

In physics, one of the possibilities to study the deviations from equilibrium is using thermodynamics, and in particular the Fluctuation Dissipation Theorem formalism^1,43–45^. A way of expressing the deviations from equilibrium within this formalism is through the relation between the linear response function and the auto-correlation function. The linear response function of an observable *x* of the system to a small enough perturbation *h* is given by

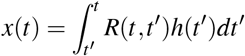

where *h*(*t*) is the perturbation that must be small. Independently, the auto-correlation (or two-point correlation function) is defined as

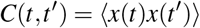

here, ⟨…⟩ stands for the average over a sufficiently large set of realizations of the object of study (i.e., an ensemble average). The formulation we are using here of the theorem states that for a system in equilibrium, the linear response function *R* is related to the auto-correlation of the unperturbed observable *x* by

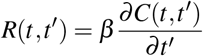

for *t*^′^ ≤ *t* and the variable *β* being identified with the inverse of the thermal bath temperature 1*/T*. For out-of-equilibrium systems, this expression is not valid and must be rewritten as

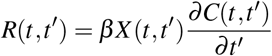

The function *X* (*t, t*^′^) is called the Fluctuation-Dissipation Ratio and characterizes the approach to equilibrium and measures the deviation from FDT. Also, a way to measure the separation from FDT, and therefore from equilibrium, is through the Integral Violation *I*(*t, t*^′^) of FDT given by

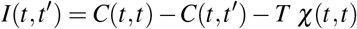

where 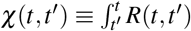*ds* is the integrated response function, or dynamic susceptibility. For systems in equilibrium, we recover the original expressions by setting *χ* = 1, *V* = 0, and *I* = 0.

Now, in the case of the system evolves with a Langevin equation

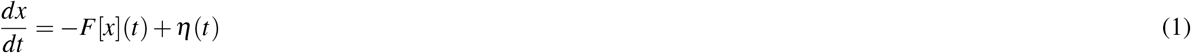

where *F* is a deterministic function and *η*(*t*) is a zero-mean Gaussian noise with auto-correlation ⟨*η*(*t*)*η*(*t*^′^)⟩ = 2*Tδ* (*t* − *t*^′^), we can reformulate the above expressions and get

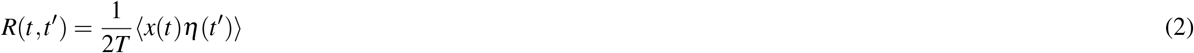

we can also define the asymmetry *A* as:

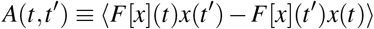

The degree of non-equilibrium could be used as a marker of brain disease states (e.g., healthy controls, mild cognitive impairment, dementia). To prove this hypothesis, we use empirical human neuroimaging data (fMRI, functional Magnetic Resonance Imaging), to write the input signals in the context of a Langevin equation, from where to obtain the linear response *R*(*t, t*^′^), which can be calculated using equation 2 further to determine *I*(*t, t*^′^). To obtain a single value for the deviation from equilibrium, it is convenient to use the *Integral Violation of the FDT* as:

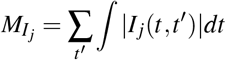

which is a metric that measures the distance from the equilibrium of the brain region *j*. Then, for each node, we take the average over all subjects as a more global indicator of distance from equilibrium for each group.

### 2.6 Model-Free Derivation

Suppose we define our observable *x*(*t*) as the BOLD signal of a given brain region. In that case, we can try to impose the Langevin equation we introduced in Equation 1 and develop the formalism presented in the previous section. The first step, to numerically compute its derivative *dx/dt*, is easy, so the only problem we face is how to split the signal into its two components, the unknown operator *F*[*x*](*t*) and the noise *η*(*t*). The key idea we use to develop the ideas further is to split the *dx/dt* signal into two components: first, we filter it by applying a Wiener filter, and we call the result − *F*[*x*](*t*) (observe the negative sign). Second, we obtain *η*(*t*) by simply adding the unfiltered and filtered versions of the signal:

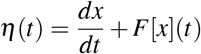

However, *η*(*t*) cannot be any function, it must satisfy two key requirements: being Gaussian, and

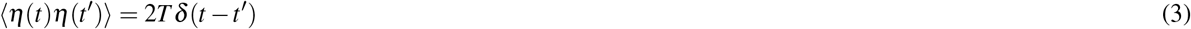

which we informally call the *sanity check* of the noise signal *η*(*t*). If the signal satisfies these conditions, we can proceed with the rest of the equilibrium analysis.

### 2.7 Model-Based Simulations of the Effects of Amyloid-Beta and Tau

This section is based on the results of our previous work^41^. In this work, we utilized the Dynamic Mean Field (DMF) model introduced by Deco et al.^36^, which employs a network model to simulate spontaneous brain activity at the whole-brain scale. According to the original formulation, each node in the model corresponds to a region of interest (i.e., a specific brain area), and the connections between these nodes represent white matter pathways. Each node, in turn, serves as a simplified representation of large populations of interconnected excitatory and inhibitory integrate-and-fire spiking neurons (with the composition of 80% excitatory and 20% inhibitory neurons, as in the original), governed by a set of dynamical equations that describe the activity of coupled excitatory (*E*) and inhibitory (*I*) neuron pools, following the reduction approach of Wong and Wang^46^. In the DMF model, excitatory synaptic currents, *I*(*E*), are mediated by NMDA receptors, while inhibitory currents, *I*(*I*), are mediated by *GABA*_*A*_ receptors. These neuronal pools are reciprocally connected, with inter-area interactions occurring exclusively at the excitatory level, scaled by the structural connectivity **C**_*k* *j*_. The DMF model can be formulated as a system of coupled differential equations:

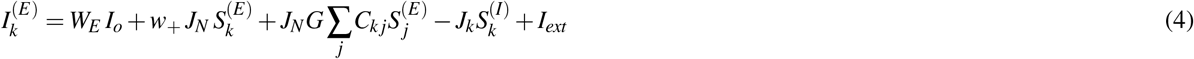

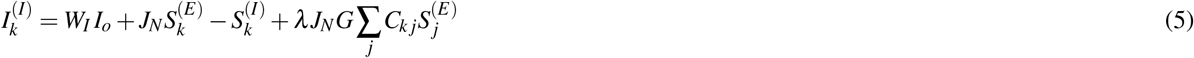

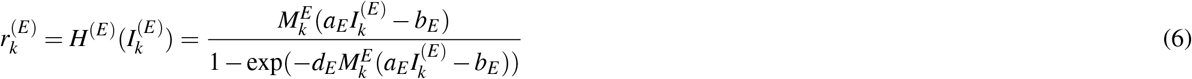

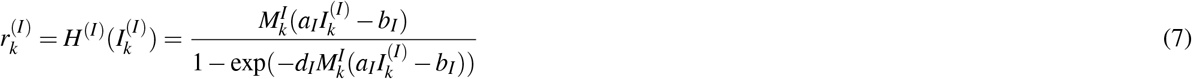

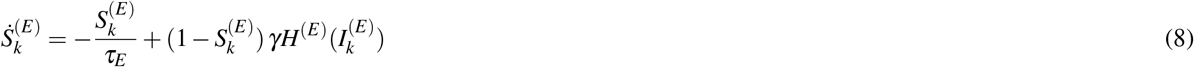

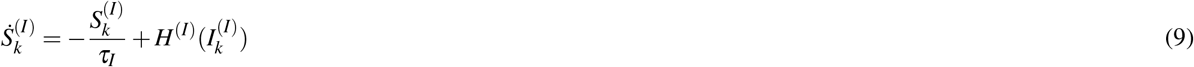

In this context, the final *two* equations should incorporate, during integration, an uncorrelated standard Gaussian noise term with an amplitude of *σ* = 0.01*nA*, utilizing the Euler-Maruyama integration method. In these equations, *λ* is a parameter that may be set to 1 or 0, depending on whether long-range feedforward inhibition is included (*λ* = 1) or excluded (*λ* = 0). In the preceding equation, the kinetic parameters are given by *γ* = 0.641*/*1000 (the factor 1000 is used to express everything in milliseconds), with *τ*_*E*_ = *τ*_*NMDA*_ and *τ*_*I*_ = *τ*_*GABA*_. The excitatory synaptic coupling is *J*_*NMDA*_ = 0.15 (nA). The overall effective external input is *I*_0_ = 0.382 (nA), scaled by *W*_*E*_ for excitatory pools and by *W*_*I*_ for inhibitory pools. The effective time constant for NMDA is *τ*_*NMDA*_ = 100 ms^46^. The values of *W*_*I*_, *I*_0_, and *J*_*NMDA*_ were selected to achieve a low level of spontaneous activity in the isolated local area model. Table 1 provides the values for the gating variables.

**Table 1.** Gating variables in the BEI model.

| Excitatory gating variables | Inhibitory gating variables |
| --- | --- |
| $a_E = 310 \text{ (nC}^{-1}\text{)}$ | $a_I = 615 \text{ (nC}^{-1}\text{)}$ |
| $b_E = 125 \text{ (Hz)}$ | $b_I = 177 \text{ (Hz)}$ |
| $d_E = 0.16 \text{ (s)}$ | $d_I = 0.087 \text{ (s)}$ |
| $\tau_E = \tau_{NMDA} = 100 \text{ (ms)}$ | $\tau_I = \tau_{GABA} = 10 \text{ (ms)}$ |
| $W_E = 1$ | $W_I = 0.7$ |

**Table 2.** Epidemiological information of the population used in this study.

| Diagnosis | n (female) | Mean age | $\sigma$ | Min. age | Max. age | Mean MMSE | $\sigma_{MMSE}$ | Min. MMSE | Max. MMSE |
| --- | --- | --- | --- | --- | --- | --- | --- | --- | --- |
| AD | 10 (5) | 72.0 | 9.6 | 55.9 | 86.1 | 21.3 | 6.8 | 9 | 30 |
| HC | 17 (10) | 70.8 | 4.3 | 63.1 | 78.0 | 29.3 | 0.7 | 28 | 30 |
| MCI | 9 (3) | 68.8 | 5.8 | 57.8 | 76.6 | 27.4 | 1.5 | 25 | 30 |

As previously mentioned, the DMF model is based on the original Wong and Wang model^46^, designed to simulate resting-state conditions. In this setup, each isolated node exhibits the characteristic noisy spontaneous activity with a low firing rate (*H*^(*E*)^ ∼3*Hz*) commonly observed in electrophysiology studies, while reusing most of the parameters from the original model. Additionally, we implemented the Feedback Inhibition Control (FIC) mechanism as described by Deco et al.^36^, where the inhibition weight *J*_*n*_ is individually adjusted for each node *n* to ensure that the firing rate of the excitatory pools *H*^(*E*)^ remains fixed at 3Hz, even when receiving excitatory input from other connected areas. Deco et al.^36^ showed that this mechanism enhances the resting-state functional connectivity (FC) prediction accuracy and yields more realistic evoked activity. We refer to this model as the balanced excitation-inhibition (BEI) model. While these local adjustments introduce regional heterogeneity, the firing rates are kept uniform across all regions, making the BEI model a homogeneous benchmark against which we can compare more advanced models that allow A*β* and tau to influence intrinsic dynamical properties across regions.

Following the Glasser parcellation^47^, we included *N* = 379 brain regions in our whole-brain network model. Each region *n* receives excitatory input into its excitatory pool from all structurally connected areas, weighted by a connectivity matrix derived from dMRI (see Section 2.7). Furthermore, all inter-regional excitatory-to-excitatory (E-to-E) connections are uniformly scaled by a global coupling factor *G*. This global scaling factor is the sole control parameter, which we adjust to bring the system to its optimal operating point, where the simulated activity best matches the empirical resting-state activity observed in healthy control participants. We conducted simulations across a range of *G* values from 0 to 5.5, with increments of 0.05 and a time step of 1 ms. For each value of *G*, we performed 200 simulations, each lasting 435 seconds, to replicate the empirical resting-state scans of 17 participants. The optimal value, identified for the *phFCD* observable, was *G* = 3.1.

After obtaining the simulated mean field activity, the next step is to convert it into a BOLD signal. For this, we employed the generalized hemodynamic model proposed by Stephan et al.^48^. The BOLD signal for the *k*-th brain region is derived from the firing rate of the excitatory pools *H*^(*E*)^. An increased firing rate leads to an increased vasodilatory signal, *s*_*k*_, which is regulated by an auto-regulatory feedback mechanism. Blood inflow *f*_*k*_ responds proportionally to this signal, resulting in changes to blood volume *v*_*k*_ and deoxyhemoglobin content *q*_*k*_. The following equations describe the relationships between these biophysical variables:

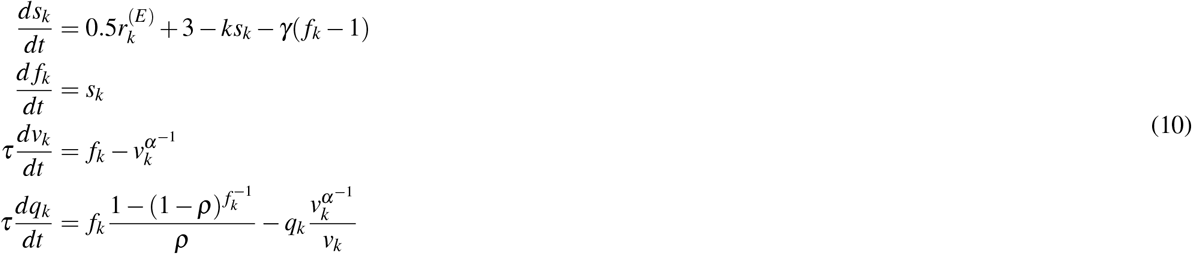

with the final measured BOLD signal being given by:

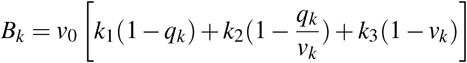

We utilized the revised version described later on^48^, which involves introducing the change of variables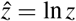. This transformation applies to the variables *z* = *f*_*k*_, *v*_*k*_, and *q*_*k*_, and alters the corresponding state equation 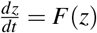 as follows:

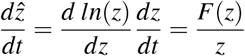

This ensures that 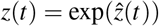 remains positive, providing appropriate support for these non-negative states and thereby enhancing numerical stability when evaluating the state equations during computation.

We introduced A*β* and Tau into our heterogeneous model at the formulae for *H*^(*E,I*)^ (the excitatory/inhibitory neuronal response functions), into the gain factor 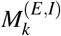 for the *k*-th area, as

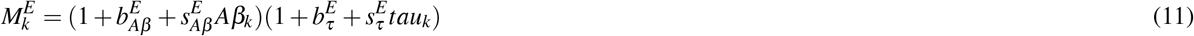

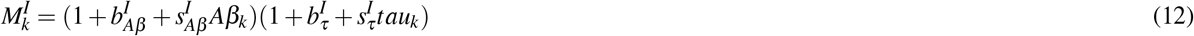

where 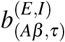 are the excitatory/inhibitory A*β* and tau bias parameters, and 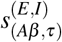 are the respective scaling factors.

We optimized these parameters subject-by-subject for our previous project and then reused the results of that previous optimization to perform the evaluations for this current project.

### 2.8 Generative Effective Connectivity Calculation

The objective of this approach is to replace the coupling matrix **C** in Equation 4 with another one 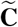 specifically tailored for each subject. To find the coupling matrix 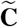, we numerically estimate it using a pseudo-gradient descent method^49, 50^. Specifically, we adjust 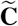 to ensure the model accurately reproduces the empirically observed covariances **FC**^empirical^ (i.e., the normalized covariance matrix from functional neuroimaging data) and the empirical time-shifted covariances **FS**^empirical^(*τ*), where *τ* represents the time lag. These covariances are normalized for each pair of regions *i* and *j* by 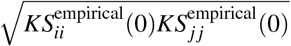. We selected the value of *τ* that resulted in a reduction of the average autocorrelation. It is important to note that fitting the time-shifted correlations can introduce asymmetries in the connectivity matrix 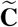, potentially leading to non-equilibrium dynamics and violations of the Fluctuation-Dissipation Theorem (FDT). These normalized time-shifted covariance matrices are obtained by dividing the shifted covariance matrix 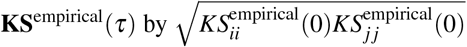 for each pair (*i, j*). Notably, these normalized time-shifted covariances disrupt the symmetry of the couplings, thereby improving the fitting accuracy^51^.

Using a heuristic pseudo-gradient algorithm, we update the 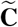 until the fit is fully optimized. More specifically, the updating uses the following form:

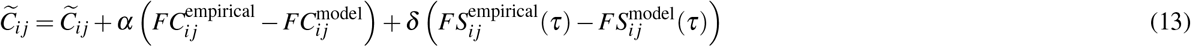

where 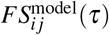 is defined similar to 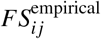. Details on the computation of the **KS** matrix can be found in the work by Deco et al.^40^. The model was run repeatedly with the updated 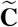 until the fit converged towards a stable value. We initialized 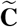 using the anatomical connectivity **C**. The **C** matrix denotes the density of fibers between cortical area *i* and *j* (obtained with probabilistic tractography from dMRI) and only updated existing connections from this matrix (in either hemisphere). We used *α* = *δ* = 1*e*^−5^ and continued until the algorithm converged. For each iteration, we compute the model results as the average over as many simulations as there are subjects. Overall, we use the term *Generative Effective Connectivity* (GEC) for the optimized 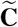^52^.

### 2.9 Comparing the Simulations with Empirical Data

As a final step for this project, we gathered the result of the DMF fitting using the procedure described in Section 2.7, as well as the GEC as described in Section 2.8, and generated the final BOLD signal for each individual with the result, thus considering the effect of both burdens and the GEC. We considered all possible combinations of both burdens and the GEC by simulating the model with the plain DMF (plus BOLD simulation) along with the standard SC, the DMF including both A*β* and tau burdens with the SC, the DMF using the GEC instead of the regular SC, and finally the DMF with both burdens and the GEC matrix. All four different simulation results were normalized such that each region’s BOLD signal had the same mean and standard deviation as the empirical data for each subject. Then the off-equilibrium FDT framework was applied to the resulting dataset.

## 3 Results

### 3.1 Model-Free analysis

The whole approach is based on the assumption that the residual noise *η*(*t*) satisfies both conditions, the Gaussianity test (*η*(*t*) must follow a Gaussian probability distribution); and the sanity check, described above. To check the first part, we exhaustively checked the resulting noise signal with several different tests, including both visual (building a histogram and comparing it with a perfect Gaussian curve, and a Q-Q plot) and analytical ones (Shapiro-Wilk’s test, D’Agostino’s *K*^2^ test, and Anderson-Darling’s test). See Figure 2A. In the case of Shapiro-Wilk’s test, 92.69% of the tests returned a positive (gaussian) result, and for D’Agostino’s *K*^2^ test, the number of positive results was 91.65%. In the case of the Anderson-Darling test, critical values provided for the normal are for the following significance levels: 15%, 10%, 5%, 2.5%, 1%, giving the following results: 15.0: 84.56%; 10.0: 90.03%; 5.0: 95.00%; 2.5: 97.39%; 1.0: 98.82%. As we can see, our analysis yielded that, in general, the vast majority of the regions have a residual noise *η*(*t*) that satisfies the Gaussianity test.

**Figure 2.**
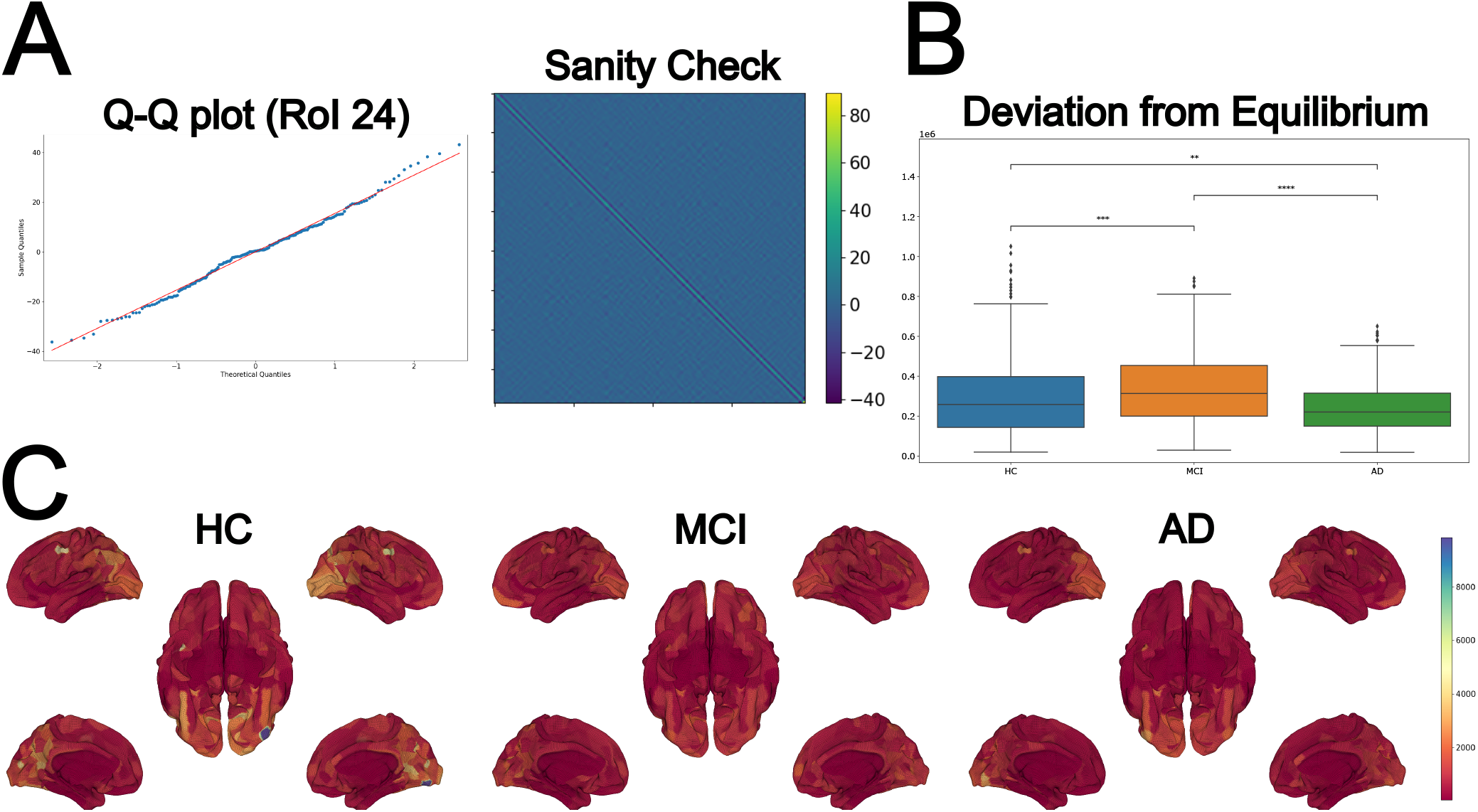
Deviation from equilibrium in Alzheimer’s Disease. **(A)** Required verifications for *η*(*t*), where on the left we can observe the Q-Q plot, which is a scatterplot created by plotting the quantiles of the derived noise against the standard normal distribution, in this case for an arbitrary area; and on the right we can observe the *sanity check*, which numerically verifies Equation 3, in this case for the whole MCI cohort. **(B)** Node-based deviation from equlibrium. Healthy controls showed lower deviations from equilibrium values across the whole-brain network than the MCI stage, reflecting the hyperactivity associated with the MCI stage, while AD shows a clear fall of the activity, showing the effect of dementia and, unfortunately, neuronal death. P-values are based on Monte–Carlo permutation tests, where **** represents *p <*= 1.00*e* − 04 and ** *p <*= 1.00*e* − 02. **(C)** Brain renders represent the deviation from equilibrium of the 379 areas, for each disease stage. We can observe that the values are higher for the healthy control cohort, and get lower and more homogeneous for subsequent stages of the disease.

Concerning the sanity check, we used Equation 3 to verify that the noise signals *η*(*t*) are adequate for our purposes, rendering the whole approach useful for neuroscience applications. We have run numerical and visual inspections, as seen in Figure 2A (right panel), to check the diagonal, i.e., for *t* = *t*^′^, Dirac delta behavior. This requirement is widely satisfied in all the tested cases, both at the subject and the node levels.

At Figure 2B we can observe that this framework allows a clean discrimination among the three cohorts, even allowing us to appreciate the increase in neuronal activity due to the effect of the first stages of the disease progression, and the subsequent fall of activity in the later ones. At Figure 2C we can see renders of the subject-based average of the node-level deviation from equilibrium coefficients. As we can see, the deviation is visible in the HC cohort. It is progressively reduced as the disease impacts the different regions along its progression, in the later stages of the disease.

Results for each RSN are presented in Figure 3, where it can be seen that strong deviations from equilibrium were minimal in the Visual network along the disease progression. Also, in its initial stages, i.e., for HC and MCI, there were no large alterations for the Dorsal Attention and the Somato-Motor networks. However, these same networks showed significant deviations for larger stages, i.e., between MCI and AD. We can observe that all other networks show an increase in their deviation from equilibrium in the first stages, followed by a pronounced decrease. This is probably due to increased neuronal excitability, as suggested by our previous results^41^.

**Figure 3.**
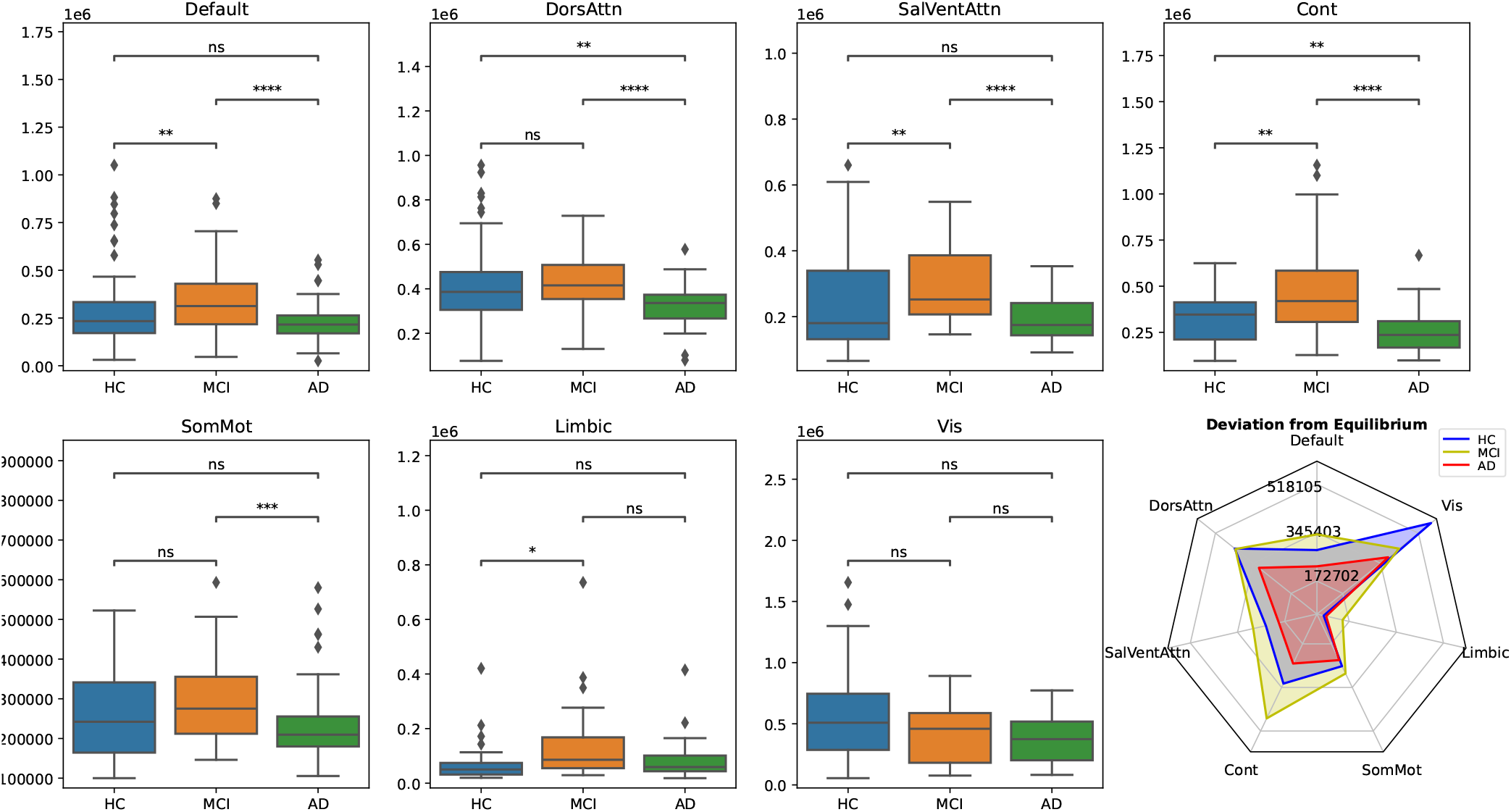
Node-based deviation from equilibrium within resting state networks. **(A)** Compared to healthy control subjects, deviation from equilibrium was preserved for the visual network at all stages. In contrast, it was preserved for Dorsal Attention and the Somato-Motor networks only between HC and MCI, but significantly decreased between the MCI and AD stages. In general, all other networks show an increase in their deviation from equilibrium in the first stages, probably due to an increase in neuronal excitability; to then show an abrupt decrease in the later stages, the result of the neuronal damage caused by the disease. p-value annotation legend: ns: *p <*= 1.00*e* + 00, *: 1.00*e* − 02 *< p <*= 5.00*e* − 02, **: 1.00*e* − 03 *< p <*= 1.00*e* − 02, ***: 1.00*e* − 04 *< p <*= 1.00*e* − 03, ****: *p <*= 1.00*e* − 04. **(B)** The radar plot represents the average deviation from equilibrium values per resting state network for each stage.

### 3.2 Model-based analysis

For the second part of this study, we compared the application of the deviation from equilibrium to the empirical data and the result of our four simulated models, see Figure 4. Namely, for each group (HC, MCI and AD), we compared the integral violation of the FDT for the DMF without burdens and with the standard SC (labeled as DMF), the DMF taking into account our previous results for the A*β* and tau burdens but with the original SC (DMF-ABeta), the DMF without burdens but computed with the GEC (DMF-GEC), and finally the result of combining both the computed burdens and the GEC (DMF-ABetaTau-GEC).

**Figure 4.**
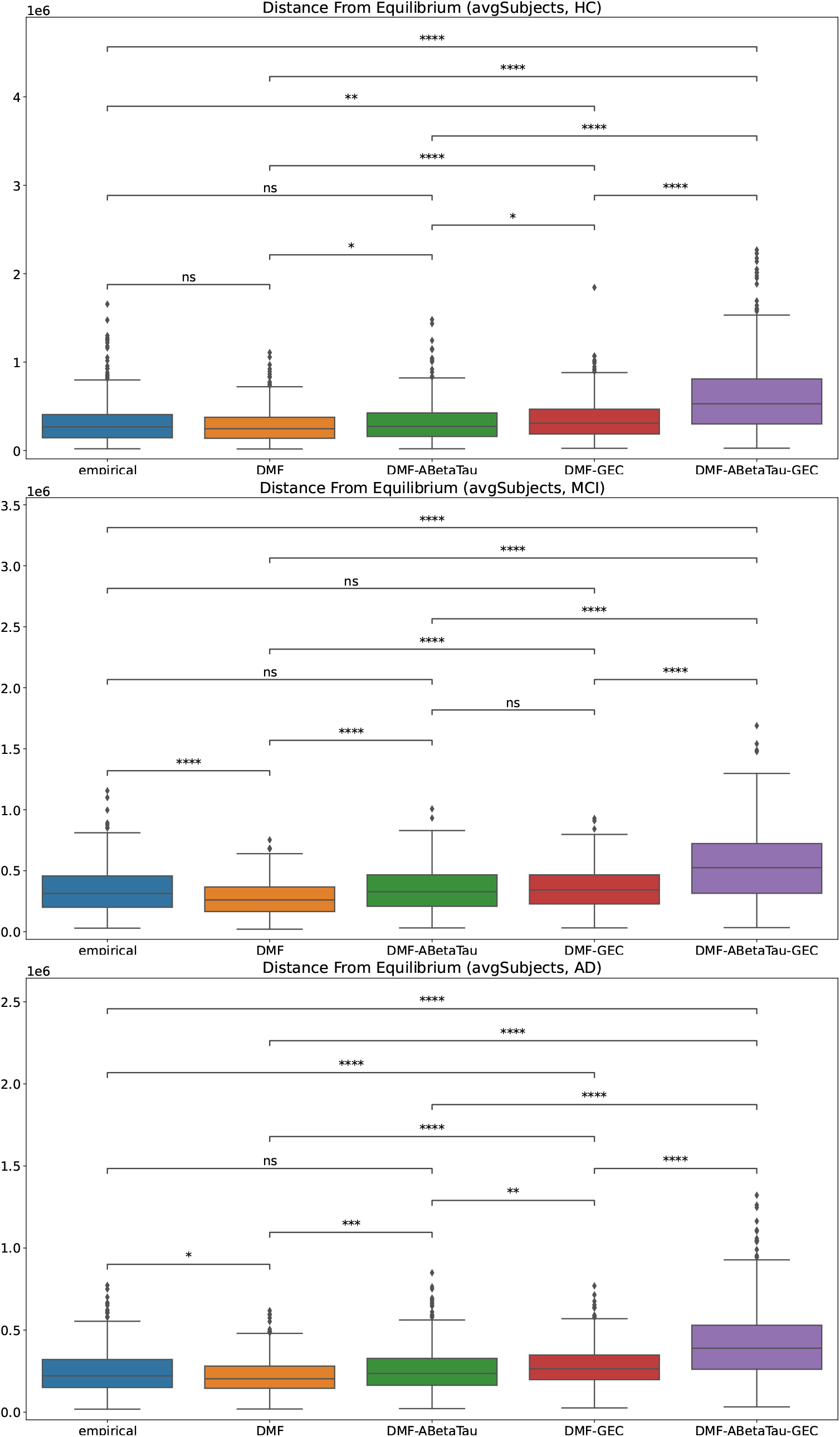
Node-based comparison of the distance from equilibrium (the integral violation of the FDT) between model-based simulations and the empirical result.

The first observation we can extract from these figures is that, in general, the off-equilibrium FDT framework can significantly distinguish between almost all combinations of empirical and simulated data, in some cases with very high sensitivity. However, it is worth noticing that, for the healthy control group, the comparison of the empirical data with the DMF model, and with the DMF model including the results for A*β* and tau, yield no significant difference, at least up to the statistical power our data allows. This is expected, as this group should not have a substantial dependence between their brain activity and the burdens related to the disease. Then, this equilibrium changes for the MCI group, as the framework has no problem distinguishing the DMF model results from the empirical data. Still, the results show it cannot differentiate between the empirical data and the simulation with the addition of A*β* and tau, or with the addition of the GEC. Also, these two simulations cannot be distinguished among themselves. Finally, for the AD group, the only non-significant comparison is the one involving the empirical data and the simulations including both burdens, evidencing the high accuracy of the mathematical model including the burdens. In this last case, the GEC can no longer compare to the empirical data, showing that long-range connections alone cannot describe the empirical results. All these comparisons can be appreciated better at Figure 5, where the different deviations from equilibrium are normalized to the empirical dataset, and their differences (in absolute value) are plotted.

**Figure 5.**
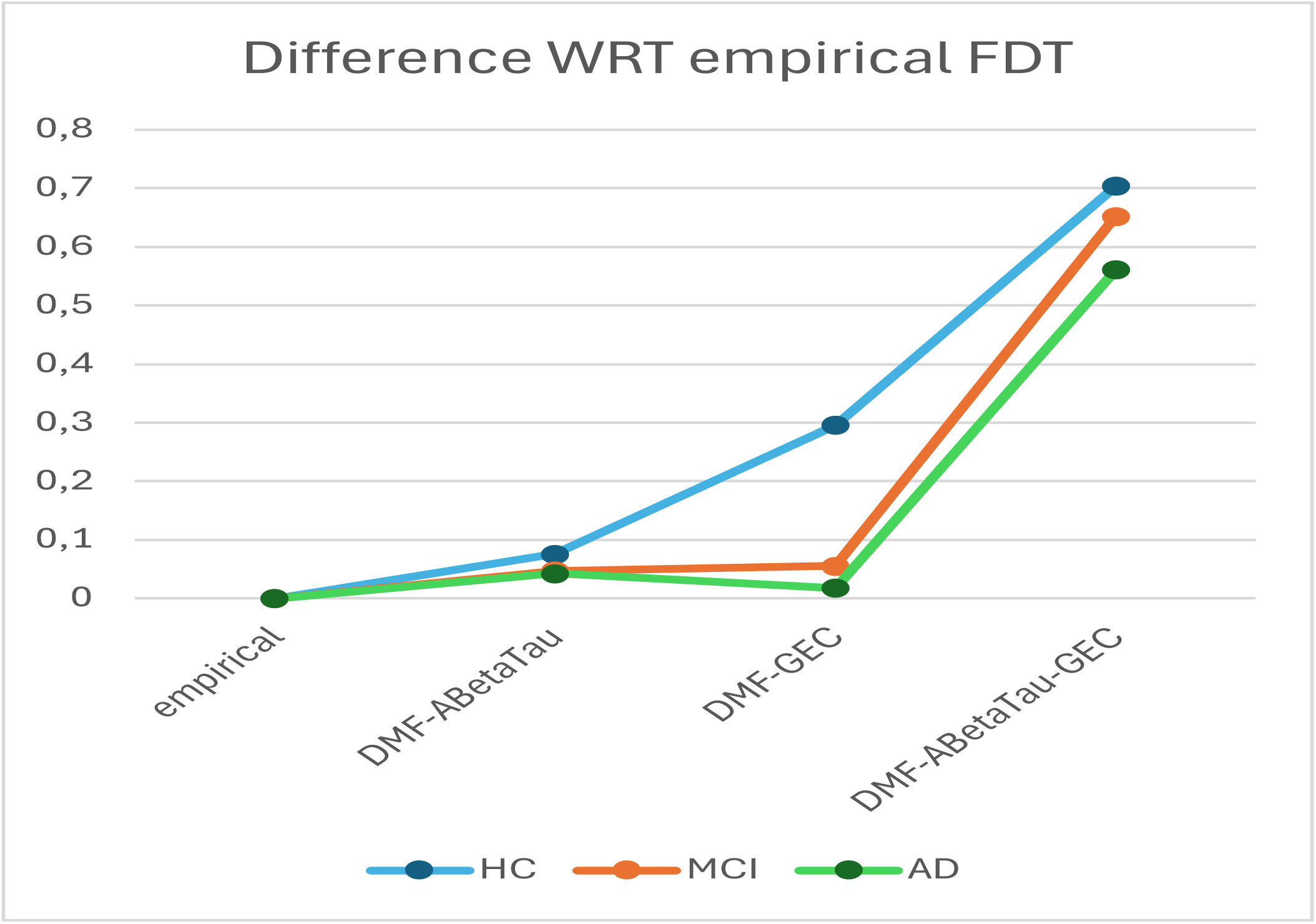
Analysis of the results in Figure 4, taking the absolute value of the difference with the corresponding empirical data as reference, which thus takes a zero value.

There is still an important piece of information we can extract from the graphs, in particular the last one: the lack of similarity between the empirical data and the results of the simulations and the combined influence of A*β*, tau, and the subject-tailored GEC matrices. This is an important result, as we discuss in the next section.

Finally, Figure 6 shows the comparison of the evaluations of the framework on the simulations of the DMF and the two burdens (i.e., A*β* and tau). As we can see, the three stages can be discerned, with the same increase in the distance from the equilibrium between HC and MCI, with the posterior decrease between MCI and AD. This same behavior was already observed in Figure 2.

**Figure 6.**
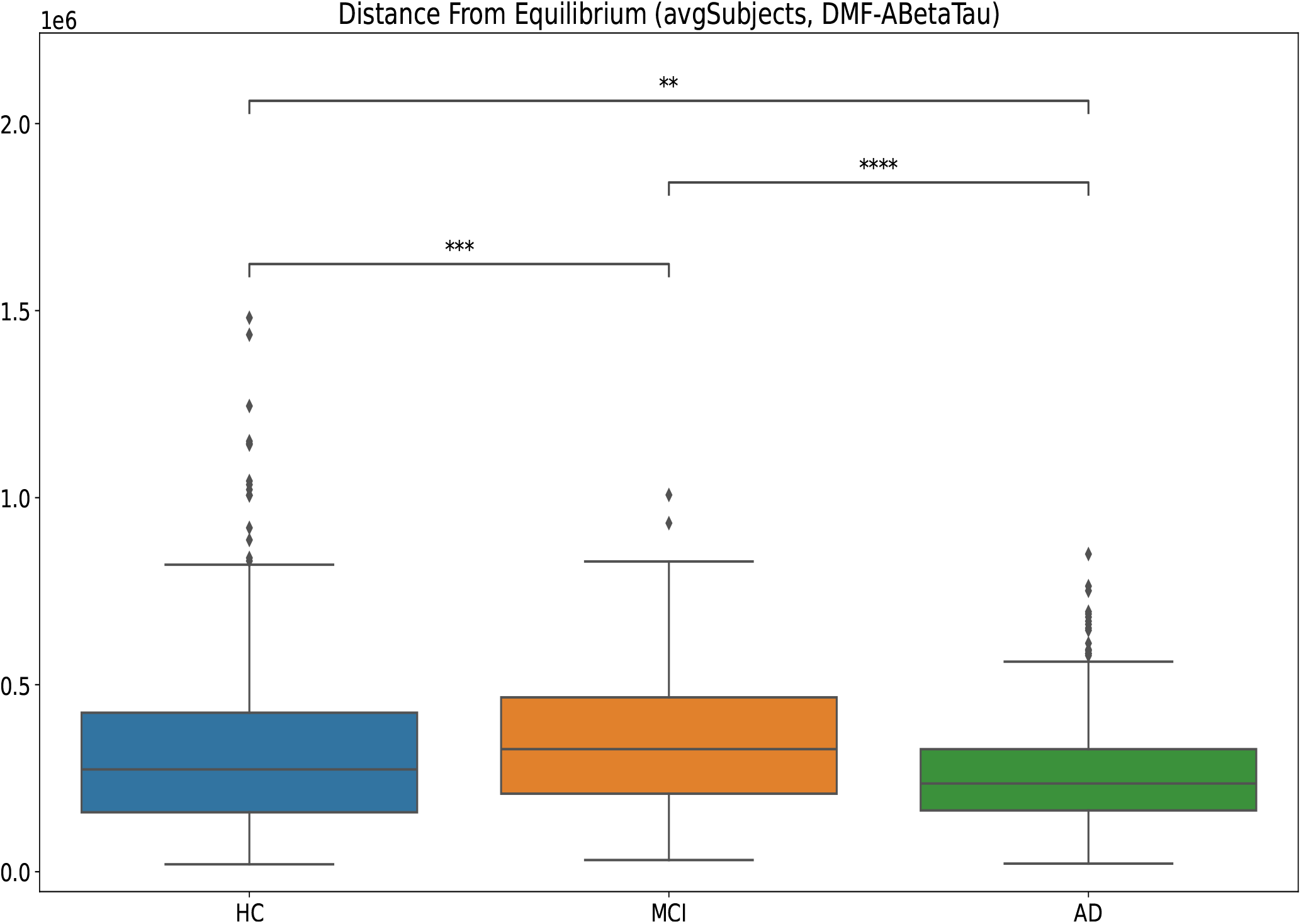
Comparison of the integral violation of the FDT for the model-based simulation taking into account the A*β* and tau burdens. We can see the same general behavior as in Figure 2, providing a mechanistic explanation of the increase-followed-by-decrease behavior in the deviations from equilibrium.

## 4 Discussion

We have presented a model-free formulation of the off-equilibrium FDT approach, which allows the processing of fMRI data directly without resorting to an intermediate simulated model. To do it, we split the input signal for each region into a base signal and noise components, checking for the latter the general verification of the required conditions imposed by the used FDT formalism, the so-called *sanity checks*. Once verified, the formalism can be safely applied to both the whole brain and the resting state network levels, resulting in an unprecedented sensitivity to detect variations in neuronal activity.

We applied this extended formalism to Alzheimer’s disease, where the data shows an increase in the deviation from equilibrium in the first stages (HC to MCI), probably because of the known hyperexcitability phenomenon in the first stages of Alzheimer’s disease; followed by a severe neuronal degradation in later stages (AD), a sign of the neuronal damage produced by the disease progression. This agrees with our previous results^41^, and can be mechanistically explained with our model-based analysis, see below.

In our RSN analysis, in general, most of the networks showed the same increase-followed-by-decrease pattern that the whole brain analysis revealed, except for the Visual network, which is well-known to be affected by the disease, but mostly in its latest stages^53,54^. Our results agree with the published findings for similar studies^53^. We can also observe that the Dorsal-Attention and the Somato-Motor networks are preserved in the first stages (MCI) but are severely affected in later stages (AD). In contrast, the Limbic network is mostly affected in the MCI stage. Altogether, we can see that the off-equilibrium FDT approach has quite good, statistically significant discerning power to identify the changes in the different RSNs, rendering them good targets as biomarkers for patient diagnosis.

All these results, both at the whole brain and at the RSN levels, should be compared with other successful frameworks, such as the Intrinsic Ignition framework^55^. In the context of this framework, the Metastability measure refers to the local degree of functional variability of each brain area over time. It has been successfully applied to the same dataset that we use here. The results show a progressive *decrease* of this measure along the disease progression, which correlates well with the sustained reduction of brain network dynamics and thus, cognitive function. the off-equilibrium FDT framework, on the other side, shows the deviations from the equilibrium of the brain activity, which probably reflects the hyperexcitability during MCI and the posterior neuronal loss in AD, see below. These results show that the off-equilibrium framework is an excellent companion to the Intrinsic Ignition framework in Alzheimer’s disease^56^, and its complementary role with other successful frameworks such as Intrinsic Ignition.

On the other hand, we compared the performance of four different simulated whole-brain models concerning the empirical data, for each group. We observed that, for the healthy control group, the simple model without any addition already does a good job explaining the empirical data. For the MCI group, adding the protein burdens, namely A*β* and tau, or using the individually fitted GEC matrices could explain the empirical results. Finally, for the AD group, only the model incorporating the local actions of A*β* and tau could mechanistically explain the data, always within the statistical power given by our data. This shows that the local affectation by the protein burdens plays a key role in disease development. Still, the long-range connections, although quite descriptive in the MCI stage, cannot completely explain the latest stages of the disease.

There is one last important result to consider, the drastic reduction in the predictive power of the mathematical model when the protein burdens are considered along with the GEC matrices. This may come as a surprise, given the high similarity (absolute value-wise) between the separate simulations. However, it also is a very enlightening result, as it shows that the local influence of the protein burdens is important, but only a part of the picture, which is far from providing a mechanistic explanation when combined with long-range connections, represented by the GEC matrices. Thus, this information gives the precious insight that more research is needed to understand the interplay between the actions of the protein burdens, A*β* and tau, and the long-range connectivity, represented by the GEC matrices.

As a final analysis, our model-based simulations can add to our understanding of the increase-followed-by-decrease behavior seen in figure 2, by plotting the results for the DMF plus A*β* and tau burdens. This result is shown in Figure 6, which presents the same *same* increase in the violation of the FDT as the empirical data. This provides a much-needed mechanistic explanation of this behavior, which can be explained by the hyperexcitability in the MCI stage, with posterior neuronal damage in the later AD stage^41^. This also provides a biologically oriented explanation for the violation of the FDT, as a measure of the activity of a brain region.

## Acknowledgements

Data collection and sharing for this project was funded by the Alzheimer’s Disease Neuroimaging Initiative (ADNI) (National Institutes of Health Grant U01 AG024904) and DOD ADNI (Department of Defense award number W81XWH-12-2-0012). ADNI is funded by the National Institute on Aging, the National Institute of Biomedical Imaging and Bioengineering, and through generous contributions from the following: AbbVie, Alzheimer’s Association; Alzheimer’s Drug Discovery Foundation; Araclon Biotech; BioClinica, Inc.; Biogen; Bristol-Myers Squibb Company; CereSpir, Inc.; Cogstate; Eisai Inc.; Elan Pharmaceuticals, Inc.; Eli Lilly and Company; EuroImmun; F. Hoffmann-La Roche Ltd and its affiliated company Genentech, Inc.; Fujirebio; GE Healthcare; IXICO Ltd.; Janssen Alzheimer Immunotherapy Research & Development, LLC.; Johnson & Johnson Pharmaceutical Research & Development LLC.; Lumosity; Lundbeck; Merck & Co., Inc.; Meso Scale Diagnostics, LLC.; NeuroRx Research; Neurotrack Technologies; Novartis Pharmaceuticals Corporation; Pfizer Inc.; Piramal Imaging; Servier; Takeda Pharmaceutical Company; and Transition Therapeutics. The Canadian Institutes of Health Research is providing funds to support ADNI clinical sites in Canada. Private sector contributions are facilitated by the Foundation for the National Institutes of Health (www.fnih.org). The grantee organization is the Northern California Institute for Research and Education, and the study is coordinated by the Alzheimer’s Therapeutic Research Institute at the University of Southern California. ADNI data are disseminated by the Laboratory for Neuro Imaging at the University of Southern California.

## Data and code availability statement

Upon acceptance, all code for implementing computational models and reproducing our results will be available at https://github.com/dagush/WholeBrain

