## Supplementary Info for "Off-Equilibrium Fluctuation-Dissipation Theorem Paves the Way in Alzheimer’s Disease Research"

### SUPPLEMENTARY MATERIAL

**Supplementary Table 1**. MPRAGE metadata.

| ID | Model | TE [ms] | TR [s] | MatrixSize | VoxelSize [mm] |
|---|---|---|---|---|---|
| 023_S_1190 | TrioTim | 2.98 | 2.3 | (176, 240, 256) | (1.0, 1.0, 1.0) |
| 002_S_1280 | Prisma_fit | 2.95 | 2.3 | (176, 240, 256) | (1.2000046, 1.0546875, 1.0546875) |
| 011_S_4547 | Prisma_fit | 2.98 | 2.3 | (208, 240, 256) | (1.0, 1.0, 1.0) |
| 168_S_6142 | Prisma_fit | 2.98 | 2.3 | (208, 240, 256) | (0.9999948, 1.0, 1.0) |
| 002_S_6103 | Prisma_fit | 2.98 | 2.3 | (208, 240, 256) | (1.0, 1.0, 1.0) |
| 002_S_4654 | Prisma_fit | 2.95 | 2.3 | (176, 240, 256) | (1.199997, 1.0546875, 1.0546875) |
| 022_S_5004 | TrioTim | 2.98 | 2.3 | (176, 240, 256) | (1.0, 1.0, 1.0) |
| 003_S_6067 | Prisma | 2.98 | 2.3 | (208, 240, 256) | (1.0, 1.0, 1.0) |
| 002_S_4229 | Prisma_fit | 2.98 | 2.3 | (208, 240, 256) | (1.0, 1.0, 1.0) |
| 012_S_6073 | Prisma | 2.98 | 2.3 | (208, 240, 256) | (1.0, 1.0, 1.0) |
| 002_S_1261 | Prisma_fit | 2.95 | 2.3 | (176, 240, 256) | (1.2000046, 1.0546875, 1.0546875) |
| 002_S_6009 | Prisma_fit | 2.95 | 2.3 | (176, 240, 256) | (1.2000046, 1.0546875, 1.0546875) |
| 007_S_4488 | Prisma | 2.98 | 2.3 | (208, 240, 256) | (1.0, 1.0, 1.0) |
| 003_S_4288 | Prisma | 2.98 | 2.3 | (208, 240, 256) | (1.0, 1.0, 1.0) |
| 002_S_4213 | Prisma_fit | 2.98 | 2.3 | (208, 240, 256) | (1.0, 1.0, 1.0) |
| 114_S_6039 | Verio | 2.98 | 2.3 | (176, 240, 256) | (1.0, 1.0, 1.0) |
| 036_S_4430 | Skyra | 2.95 | 2.3 | (176, 240, 256) | (1.199997, 1.0546875, 1.0546875) |
| 041_S_4974 | Prisma_fit | 2.95 | 2.3 | (176, 240, 256) | (1.2000046, 1.0546875, 1.0546875) |
| 007_S_4272 | Prisma | 2.98 | 2.3 | (208, 240, 256) | (1.0, 1.0, 1.0) |
| 011_S_4827 | Prisma_fit | 2.98 | 2.3 | (208, 240, 256) | (1.0, 1.0, 1.0) |
| 002_S_6053 | Prisma_fit | 2.98 | 2.3 | (208, 240, 256) | (1.0, 1.0, 1.0) |
| 003_S_4644 | Prisma | 2.98 | 2.3 | (208, 240, 256) | (1.0, 1.0, 1.0) |
| 002_S_4799 | Prisma_fit | 2.95 | 2.3 | (176, 240, 256) | (1.2000046, 1.0546875, 1.0546875) |
| 002_S_0413 | Prisma_fit | 2.95 | 2.3 | (176, 240, 256) | (1.2000046, 1.0546875, 1.0546875) |
| 114_S_0416 | Verio | 2.98 | 2.3 | (176, 240, 256) | (1.0, 1.0, 1.0) |
| 002_S_5178 | Prisma_fit | 2.95 | 2.3 | (176, 240, 256) | (1.199997, 1.0546875, 1.0546875) |
| 002_S_6030 | Prisma_fit | 2.95 | 2.3 | (176, 240, 256) | (1.2000046, 1.0546875, 1.0546875) |
| 003_S_1122 | Prisma | 2.98 | 2.3 | (208, 240, 256) | (1.0000056, 1.0, 1.0) |

| | | | | | |
|---|---|---|---|---|---|
| 011_S_4893 | Prisma_fit | 2.98 | 2.3 | (208, 240, 256) | (1.0, 1.0, 1.0) |
| 002_S_1155 | Prisma_fit | 2.95 | 2.3 | (176, 240, 256) | (1.2000046, 1.0546875, 1.0546875) |
| 036_S_4715 | Skyra | 2.95 | 2.3 | (176, 240, 256) | (1.2000046, 1.0546875, 1.0546875) |
| 007_S_4387 | Prisma | 2.98 | 2.3 | (208, 240, 256) | (1.0, 1.0, 1.0) |
| 007_S_4620 | Prisma | 2.98 | 2.3 | (208, 240, 256) | (1.0, 1.0, 1.0) |

**Supplementary Table 2**. FLAIR metadata.

| ID | Model | TE [ms] | TR [s] | MatrixSize | VoxelSize [mm] |
|---|---|---|---|---|---|
| 023_S_1190 | TrioTim | 443 | 4.8 | (160, 256, 256) | (1.2000046, 1.0, 1.0) |
| 002_S_1280 | Prisma_fit | 441 | 4.8 | (160, 256, 256) | (1.2000046, 1.0, 1.0) |
| 011_S_4547 | Prisma_fit | 441 | 4.8 | (160, 256, 256) | (1.2000046, 1.0, 1.0) |
| 168_S_6142 | Prisma_fit | 441 | 4.8 | (160, 256, 256) | (1.2000002, 1.0, 1.0) |
| 002_S_6103 | Prisma_fit | 441 | 4.8 | (160, 256, 256) | (1.199997, 1.0, 1.0) |
| 002_S_4654 | Prisma_fit | 441 | 4.8 | (160, 256, 256) | (1.199997, 1.0, 1.0) |
| 022_S_5004 | TrioTim | 439 | 4.8 | (160, 256, 256) | (1.199997, 1.0, 1.0) |
| 003_S_6067 | Prisma | 441 | 4.8 | (160, 256, 256) | (1.2000046, 1.0, 1.0) |
| 002_S_4229 | Prisma_fit | 441 | 4.8 | (160, 256, 256) | (1.199997, 1.0, 1.0) |
| 012_S_6073 | Prisma | 441 | 4.8 | (160, 256, 256) | (1.2000046, 1.0, 1.0) |
| 002_S_1261 | Prisma_fit | 441 | 4.8 | (160, 256, 256) | (1.2000046, 1.0, 1.0) |
| 002_S_6009 | Prisma_fit | 441 | 4.8 | (160, 256, 256) | (1.2000046, 1.0, 1.0) |
| 007_S_4488 | Prisma | 441 | 4.8 | (160, 256, 256) | (1.2000046, 1.0, 1.0) |
| 003_S_4288 | Prisma | 441 | 4.8 | (160, 256, 256) | (1.199997, 1.0, 1.0) |
| 002_S_4213 | Prisma_fit | 441 | 4.8 | (160, 256, 256) | (1.199997, 1.0, 1.0) |
| 114_S_6039 | Verio | 343 | 4.8 | (160, 256, 256) | (1.0, 1.0, 1.0) |
| 036_S_4430 | Skyra | 441 | 4.8 | (160, 256, 256) | (1.2000046, 1.0, 1.0) |
| 041_S_4974 | Prisma_fit | 441 | 4.8 | (160, 256, 256) | (1.199997, 1.0, 1.0) |
| 007_S_4272 | Prisma | 441 | 4.8 | (160, 256, 256) | (1.199997, 1.0, 1.0) |
| 011_S_4827 | Prisma_fit | 441 | 4.8 | (160, 256, 256) | (1.199997, 1.0, 1.0) |
| 002_S_6053 | Prisma_fit | 441 | 4.8 | (160, 256, 256) | (1.199997, 1.0, 1.0) |
| 003_S_4644 | Prisma | 441 | 4.8 | (160, 256, 256) | (1.199997, 1.0, 1.0) |
| 002_S_4799 | Prisma_fit | 441 | 4.8 | (160, 256, 256) | (1.2000046, 1.0, 1.0) |
| 002_S_0413 | Prisma_fit | 441 | 4.8 | (160, 256, 256) | (1.2000046, 1.0, 1.0) |
| 114_S_0416 | Verio | 343 | 4.8 | (160, 256, 256) | (1.0, 1.0, 1.0) |
| 002_S_5178 | Prisma_fit | 441 | 4.8 | (160, 256, 256) | (1.199997, 1.0, 1.0) |
| 002_S_6030 | Prisma_fit | 441 | 4.8 | (160, 256, 256) | (1.2000046, 1.0, 1.0) |
| 003_S_1122 | Prisma | 441 | 4.8 | (160, 256, 256) | (1.2000005, 1.0, 1.0) |
| 011_S_4893 | Prisma_fit | 441 | 4.8 | (160, 256, 256) | (1.2000046, 1.0, 1.0) |
| 002_S_1155 | Prisma_fit | 441 | 4.8 | (160, 256, 256) | (1.2000046, 1.0, 1.0) |
| 036_S_4715 | Skyra | 441 | 4.8 | (160, 256, 256) | (1.2000046, 1.0, 1.0) |
| 007_S_4387 | Prisma | 441 | 4.8 | (160, 256, 256) | (1.2000046, 1.0, 1.0) |
| 007_S_4620 | Prisma | 441 | 4.8 | (160, 256, 256) | (1.2000046, 1.0, 1.0) |

**Supplementary Table 3**. DTI metadata (only for HC participants to average the SC template)

| ID | Model | Institute | TE [ms] | TR [s] | MatrixSize | VoxelSize [mm, mm, mm, s] | n_Bvec s | Bvals |
|---|---|---|---|---|---|---|---|---|
| 002_S_1 280 | Prisma_ fit | OHSU_AIRC | 56 | 7.2 | (116, 116, 80,55) | (2.0, 2.0, 2.0, 7.2) | 49 | [ 0. 1000.] |
| 002_S_6 103 | Prisma_ fit | OHSU_AIRC | 56 | 7.2 | (116, 116, 80,55) | (2.0, 2.0, 2.0, 7.2) | 49 | [ 0. 1000.] |
| 003_S_6 067 | Prisma | USCINI | 56 | 7.2 | (116, 116, 80,55) | (2.0, 2.0, 2.0, 7.2) | 49 | [ 0. 1000.] |
| 002_S_6 009 | Prisma_ fit | OHSU_AIRC | 56 | 7.2 | (116, 116, 80,55) | (2.0, 2.0, 2.0, 7.2) | 49 | [ 0. 1000.] |
| 007_S_4 488 | Prisma | MAYO_CLINIC_MR I_58 | 71 | 3.4 | (116, 116, 81,127) | (2.0, 2.0, 2.0, 3.4) | 115 | [ 0. 500. 1000. 2000.] |
| 003_S_4 288 | Prisma | USC_Stevens_Hall_I nstitut e | 56 | 7.2 | (116, 116, 80,55) | (2.0, 2.0, 2.0, 7.2) | 49 | [ 0. 1000.] |
| 002_S_4 213 | Prisma_ fit | OHSU_AIRC | 56 | 7.2 | (116, 116, 80,55) | (2.5172415, 2.5172415, 2.0, 7.2) | 49 | [ 0. 1000.] |
| 002_S_6 053 | Prisma_ fit | OHSU_AIRC | 56 | 7.2 | (116, 116, 80,55) | (2.0, 2.0, 2.0, 7.2) | 49 | [ 0. 1000.] |
| 003_S_4 644 | Prisma | USCINI | 56 | 7.2 | (116, 116, 80,55) | (2.0, 2.0, 2.0, 7.2) | 49 | [ 0. 1000.] |
| 002_S_4 799 | Prisma_ fit | OHSU_AIRC | 56 | 7.2 | (116, 116, 80,55) | (2.0, 2.0, 2.0, 7.2) | 49 | [ 0. 1000.] |
| 002_S_0 413 | Prisma_ fit | OHSU_AIRC | 56 | 7.2 | (116, 116, 80,55) | (2.0, 2.0, 2.0, 7.2) | 49 | [ 0. 1000.] |
| 002_S_5 178 | Prisma_ fit | OHSU_AIRC | 56 | 7.2 | (116, 116, 80,55) | (2.0, 2.0, 2.0, 7.2) | 49 | [ 0. 1000.] |
| 002_S_6 030 | Prisma_ fit | OHSU_AIRC | 56 | 7.2 | (116, 116, 80,55) | (2.0, 2.0, 2.0, 7.2) | 49 | [ 0. 1000.] |
| 007_S_4 387 | Prisma | MAYO_CLINIC_MR I_58 | 71 | 3.4 | (116, 116, 81,127) | (2.0, 2.0, 2.0, 3.4) | 115 | [ 0. 500. 1000. 2000.] |
| 007_S_4 620 | Prisma | MAYO_CLINIC_MR I_58 | 71 | 3.4 | (116, 116, 81,127) | (2.0, 2.0, 2.0, 3.4) | 115 | [ 0. 500. 1000. 2000.] |

**Supplementary Table 4.** AV-45 PET (Abeta) metadata.

| ID | Scanner | Model | MatrixSize | VoxelSize [mm] |
|---|---|---|---|---|
| 023_S_1190 | Siemens | Biograph6_TruePoint | (160, 160, 96) | (1.5, 1.5, 1.5) |
| 002_S_1280 | Philips | GEMINI_TF_TOF_16 | (160, 160, 96) | (1.5, 1.5, 1.5) |
| 011_S_4547 | Siemens | Biograph40_TruePoint | (160, 160, 96) | (1.5, 1.5, 1.5) |
| 168_S_6142 | GE | Discovery_STE | (160, 160, 96) | (1.5, 1.5, 1.5) |
| 002_S_6103 | Philips | GEMINI_TF_TOF_16 | (160, 160, 96) | (1.5, 1.5, 1.5) |
| 002_S_4654 | Philips | GEMINI_TF_TOF_16 | (160, 160, 96) | (1.5, 1.5, 1.5) |
| 022_S_5004 | Philips | Ingenuity_TF_PET_CT | (160, 160, 96) | (1.5, 1.5, 1.5) |
| 003_S_6067 | Siemens | Biograph64_TruePoint | (160, 160, 96) | (1.5, 1.5, 1.5) |
| 002_S_4229 | Philips | GEMINI_TF_TOF_16 | (160, 160, 96) | (1.5, 1.5, 1.5) |
| 012_S_6073 | GE | Discovery_710 | (160, 160, 96) | (1.5, 1.5, 1.5) |
| 002_S_1261 | Philips | GEMINI_TF_TOF_16 | (160, 160, 96) | (1.5, 1.5, 1.5) |
| 002_S_6009 | Philips | GEMINI_TF_TOF_16 | (160, 160, 96) | (1.5, 1.5, 1.5) |
| 007_S_4488 | GE | Discovery_690 | (160, 160, 96) | (1.5, 1.5, 1.5) |
| 003_S_4288 | Siemens | Biograph64_TruePoint | (160, 160, 96) | (1.5, 1.5, 1.5) |
| 002_S_4213 | Philips | GEMINI_TF_TOF_16 | (160, 160, 96) | (1.5, 1.5, 1.5) |
| 114_S_6039 | Philips | GEMINI_TF_TOF_64 | (160, 160, 96) | (1.5, 1.5, 1.5) |
| 036_S_4430 | Siemens | Biograph40_TruePoint | (160, 160, 96) | (1.5, 1.5, 1.5) |
| 041_S_4974 | Siemens | HR+ | (160, 160, 96) | (1.5, 1.5, 1.5) |
| 007_S_4272 | GE | Discovery_690 | (160, 160, 96) | (1.5, 1.5, 1.5) |
| 011_S_4827 | Siemens | Biograph40_TruePoint | (160, 160, 96) | (1.5, 1.5, 1.5) |
| 002_S_6053 | Philips | GEMINI_TF_TOF_16 | (160, 160, 96) | (1.5, 1.5, 1.5) |
| 003_S_4644 | Siemens | Biograph64_TruePoint | (160, 160, 96) | (1.5, 1.5, 1.5) |
| 002_S_4799 | Philips | GEMINI_TF_TOF_16 | (160, 160, 96) | (1.5, 1.5, 1.5) |
| 002_S_0413 | Philips | GEMINI_TF_TOF_16 | (160, 160, 96) | (1.5, 1.5, 1.5) |
| 114_S_0416 | Philips | GEMINI_TF_TOF_64 | (160, 160, 96) | (1.5, 1.5, 1.5) |
| 002_S_5178 | Philips | GEMINI_TF_TOF_16 | (160, 160, 96) | (1.5, 1.5, 1.5) |
| 002_S_6030 | Philips | GEMINI_TF_TOF_16 | (160, 160, 96) | (1.5, 1.5, 1.5) |
| 003_S_1122 | Siemens | Biograph64_TruePoint | (160, 160, 96) | (1.5, 1.5, 1.5) |
| 011_S_4893 | Siemens | Biograph40_TruePoint | (160, 160, 96) | (1.5, 1.5, 1.5) |
| 002_S_1155 | Philips | GEMINI_TF_TOF_16 | (160, 160, 96) | (1.5, 1.5, 1.5) |
| 036_S_4715 | Siemens | Biograph40_TruePoint | (160, 160, 96) | (1.5, 1.5, 1.5) |
| 007_S_4387 | GE | Discovery_690 | (160, 160, 96) | (1.5, 1.5, 1.5) |
| 007_S_4620 | GE | Discovery_690 | (160, 160, 96) | (1.5, 1.5, 1.5) |

**Supplementary Table 5.** AV-14-51 PET (Tau) metadata

| ID | Scanner | Model | MatrixSize | VoxelSize [mm] |
|---|---|---|---|---|
| 023_S_1190 | Siemens | Biograph6_TruePoint | (160, 160, 96) | (1.5, 1.5, 1.5) |
| 002_S_1280 | Philips | GEMINI_TF_TOF_16 | (160, 160, 96) | (1.5, 1.5, 1.5) |
| 011_S_4547 | Siemens | Biograph40_TruePoint | (160, 160, 96) | (1.5, 1.5, 1.5) |
| 168_S_6142 | GE | Discovery_STE | (160, 160, 96) | (1.5, 1.5, 1.5) |
| 002_S_6103 | Philips | GEMINI_TF_TOF_16 | (160, 160, 96) | (1.5, 1.5, 1.5) |
| 002_S_4654 | Philips | GEMINI_TF_TOF_16 | (160, 160, 96) | (1.5, 1.5, 1.5) |
| 022_S_5004 | Philips | Ingenuity_TF_PET_CT | (160, 160, 96) | (1.5, 1.5, 1.5) |
| 003_S_6067 | Siemens | Biograph64_TruePoint | (160, 160, 96) | (1.5, 1.5, 1.5) |
| 002_S_4229 | Philips | GEMINI_TF_TOF_16 | (160, 160, 96) | (1.5, 1.5, 1.5) |
| 012_S_6073 | GE | Discovery_710 | (160, 160, 96) | (1.5, 1.5, 1.5) |
| 002_S_1261 | Philips | GEMINI_TF_TOF_16 | (160, 160, 96) | (1.5, 1.5, 1.5) |
| 002_S_6009 | Philips | GEMINI_TF_TOF_16 | (160, 160, 96) | (1.5, 1.5, 1.5) |
| 007_S_4488 | GE | Discovery_690 | (160, 160, 96) | (1.5, 1.5, 1.5) |
| 003_S_4288 | Siemens | Biograph64_TruePoint | (160, 160, 96) | (1.5, 1.5, 1.5) |
| 002_S_4213 | Philips | GEMINI_TF_TOF_16 | (160, 160, 96) | (1.5, 1.5, 1.5) |
| 114_S_6039 | Philips | GEMINI_TF_TOF_64 | (160, 160, 96) | (1.5, 1.5, 1.5) |
| 036_S_4430 | Siemens | Biograph40_TruePoint | (160, 160, 96) | (1.5, 1.5, 1.5) |
| 041_S_4974 | Siemens | HR+ | (160, 160, 96) | (1.5, 1.5, 1.5) |
| 007_S_4272 | GE | Discovery_690 | (160, 160, 96) | (1.5, 1.5, 1.5) |
| 011_S_4827 | Siemens | Biograph40_TruePoint | (160, 160, 96) | (1.5, 1.5, 1.5) |
| 002_S_6053 | Philips | GEMINI_TF_TOF_16 | (160, 160, 96) | (1.5, 1.5, 1.5) |
| 003_S_4644 | Siemens | Biograph64_TruePoint | (160, 160, 96) | (1.5, 1.5, 1.5) |
| 002_S_4799 | Philips | GEMINI_TF_TOF_16 | (160, 160, 96) | (1.5, 1.5, 1.5) |
| 002_S_0413 | Philips | GEMINI_TF_TOF_16 | (160, 160, 96) | (1.5, 1.5, 1.5) |
| 114_S_0416 | Philips | GEMINI_TF_TOF_64 | (160, 160, 96) | (1.5, 1.5, 1.5) |
| 002_S_5178 | Philips | GEMINI_TF_TOF_16 | (160, 160, 96) | (1.5, 1.5, 1.5) |
| 002_S_6030 | Philips | GEMINI_TF_TOF_16 | (160, 160, 96) | (1.5, 1.5, 1.5) |
| 003_S_1122 | Siemens | Biograph64_TruePoint | (160, 160, 96) | (1.5, 1.5, 1.5) |
| 011_S_4893 | Siemens | Biograph40_TruePoint | (160, 160, 96) | (1.5, 1.5, 1.5) |
| 002_S_1155 | Philips | GEMINI_TF_TOF_16 | (160, 160, 96) | (1.5, 1.5, 1.5) |
| 036_S_4715 | Siemens | Biograph40_TruePoint | (160, 160, 96) | (1.5, 1.5, 1.5) |
| 007_S_4387 | GE | Discovery_690 | (160, 160, 96) | (1.5, 1.5, 1.5) |
| 007_S_4620 | GE | Discovery_690 | (160, 160, 96) | (1.5, 1.5, 1.5) |

**Supplementary Table 6.** Dates of Imaging and MMSE.

| ID | MMSE date | MPRAGE date | FLAIR date | DTI date | AV-45 PET date | AV-1451 PET date |
|---|---|---|---|---|---|---|
| 023_S_1190 | 17/11/13 | 17/10/23 | 17/10/23 | | 17/10/25 | 17/11/08 |
| 002_S_1280 | 18/3/7 | 17/3/13 | 17/3/13 | 17/3/13 | 17/3/02 | 18/3/5 |
| 011_S_4547 | 17/8/18 | 17/8/18 | 17/8/18 | | 17/8/30 | 17/8/24 |
| 168_S_6142 | 17/12/5 | 17/12/18 | 17/12/18 | | 18/1/17 | 18/1/3 |
| 002_S_6103 | 17/10/25 | 17/11/20 | 17/11/20 | 17/11/20 | 17/11/21 | 18/1/17 |
| 002_S_4654 | 18/5/15 | 17/5/3 | 17/5/3 | | 17/5/2 | 18/5/22 |
| 022_S_5004 | 18/6/29 | 18/3/14 | 17/3/21 | | 17/3/21 | 17/4/5 |
| 003_S_6067 | 17/12/4 | 17/8/18 | 17/8/18 | 17/8/18 | 17/10/13 | 17/10/18 |
| 002_S_4229 | 18/5/14 | 17/9/20 | 17/9/20 | | 17/9/20 | 17/10/3 |
| 012_S_6073 | 17/9/18 | 17/9/22 | 17/9/22 | | 17/10/12 | 17/10/11 |
| 002_S_1261 | 18/3/8 | 17/3/15 | 17/3/15 | | 17/3/14 | 17/3/15 |
| 002_S_6009 | 17/4/1 | 17/4/17 | 17/4/17 | 17/4/17 | 17/5/16 | 17/5/15 |
| 007_S_4488 | 18/6/11 | 17/9/12 | 17/9/12 | 17/9/12 | 17/9/22 | 17/9/13 |
| 003_S_4288 | 17/10/2 | 17/10/3 | 17/10/3 | 17/10/3 | 17/10/3 | 18/2/22 |
| 002_S_4213 | 17/8/16 | 17/8/14 | 17/8/14 | 17/8/14 | 17/8/14 | 17/8/17 |
| 114_S_6039 | 17/8/10 | 17/7/21 | 17/7/21 | | 17/8/24 | 17/10/4 |
| 036_S_4430 | 17/11/15 | 17/11/07 | 17/11/07 | | 17/11/15 | 17/11/21 |
| 041_S_4974 | 17/10/30 | 17/10/5 | 17/10/5 | | 17/8/24 | 17/10/12 |
| 007_S_4272 | 18/1/18 | 18/1/16 | 18/1/16 | | 17/12/19 | 18/1/17 |
| 011_S_4827 | 17/8/24 | 17/8/31 | 17/8/31 | | 17/8/28 | 17/9/7 |
| 002_S_6053 | 17/7/21 | 17/7/18 | 17/7/18 | 17/7/18 | 17/8/23 | 17/8/24 |
| 003_S_4644 | 17/6/26 | 17/6/21 | 17/6/21 | 17/6/21 | 18/2/28 | 18/4/17 |
| 002_S_4799 | 18/6/7 | 17/5/22 | 17/5/22 | 17/5/22 | 17/5/18 | 18/6/13 |
| 002_S_0413 | 17/6/16 | 17/6/21 | 17/6/21 | 17/6/21 | 17/6/15 | 17/6/21 |
| 114_S_0416 | 18/7/24 | 17/10/24 | 17/10/24 | | 17/10/24 | 17/11/21 |
| 002_S_5178 | 17/6/6 | 17/5/31 | 17/5/31 | 17/5/31 | 17/6/5 | 17/5/31 |
| 002_S_6030 | 17/6/9 | 17/6/15 | 17/6/15 | 17/6/15 | 17/7/25 | 17/7/24 |
| 003_S_1122 | 18/7/25 | 17/5/18 | 17/5/18 | | 17/8/8 | 17/8/10 |
| 011_S_4893 | 18/7/17 | 17/11/8 | 17/11/8 | | 17/11/1 | 17/11/7 |
| 002_S_1155 | 18/5/9 | 17/4/24 | 17/4/24 | | 17/4/20 | 17/4/24 |
| 036_S_4715 | 17/10/13 | 17/10/10 | 17/10/10 | | 17/10/10 | 17/10/12 |
| 007_S_4387 | 17/10/31 | 17/11/1 | 17/11/1 | 17/11/1 | 17/10/24 | 17/11/29 |
| 007_S_4620 | 17/12/12 | 17/12/05 | 17/12/05 | 17/12/05 | 17/12/06 | 17/12/14 |
